# Arterial mechanics, extracellular matrix, and smooth muscle differentiation in carotid arteries deficient for Rac1

**DOI:** 10.1101/2023.11.15.567271

**Authors:** Richard K. Assoian, Tina Xu, Emilia Roberts

## Abstract

Stiffening of the extracellular matrix (ECM) occurs after vascular injury and contributes to the injury-associated proliferation of vascular smooth muscle cells (SMCs). ECM stiffness also activates Rac-GTP, and SMC Rac1 deletion strongly reduces the proliferative response to injury *in vivo*. However, ECM stiffening and Rac can affect SMC differentiation, which, in itself, can influence ECM stiffness and proliferation. Here, we used pressure myography and immunofluorescence analysis of mouse carotid arteries to ask if the reported effect of Rac1 deletion on *in vivo* SMC proliferation might be secondary to a Rac effect on basal arterial stiffness or SMC differentiation. The results show that Rac1 deletion does not affect the abundance of arterial collagen-I, -III, or -V, the integrity of arterial elastin, or the arterial responses to pressure, including the axial and circumferential stretch-strain relationships that are assessments of arterial stiffness. Medial abundance of alpha-smooth muscle actin and smooth muscle-myosin heavy chain, markers of the SMC differentiated phenotype, were not statistically different in carotid arteries containing or deficient in Rac1. Nor did Rac1 deficiency have a statistically significant effect on carotid artery contraction to KCl. Overall, these data argue that the inhibitory effect of Rac1 deletion on *in vivo* SMC proliferation reflects a primary effect of Rac1 signaling to the cell cycle rather than a secondary effect associated with altered SMC differentiation or arterial stiffness.

## Introduction

Large elastic and muscular arteries such as the carotid and femoral contain a single-cell layer of endothelial cells (called the intima) that provides a non-thrombogenic surface around the lumen, a thick layer of vascular smooth muscle cells (SMCs) (called the “media”), and a diffuse outer layer (called the “adventitia”) that typically contains fibroblasts, inflammatory, and precursor cells [1–4].

Vascular SMCs are highly plastic and can undergo reversible differentiation-dedifferentiation [5–8]. *In vivo*, arterial SMCs normally exist in the differentiated “contractile” state characterized by periodic contraction, quiescence, and high expression of contractile proteins such as smooth muscle myosin heavy chain (SM-MHC) and alpha-smooth muscle actin (α-SMA). However, SMC dedifferentiation to a “synthetic” phenotype is a hallmark of several vascular pathologies including atherosclerosis and the response to vascular injury. This synthetic phenotype is characterized by reduced contraction and an increased capacity to proliferate and migrate. Additionally, remodeling of the extracellular matrix (ECM) occurs during SMC dedifferentiation to the synthetic phenotype and is thought to contribute to the arterial stiffening seen after vascular injury and in atherosclerosis [9, 10]. Arterial stiffening, in turn, is permissive for cell proliferation and migration [9, 11]. While the arterial ECM is highly complex, several studies have shown that two of the best studied components, elastin and the fibrillar collagens, have distinguishable roles in arterial mechanics. Elastin allows for arterial recoil and dominates arterial stiffness at low stress and stretch while the fibrillar collagens regulate arterial stiffness at higher stress and stretch [2]. Altered expression, degradation and crosslinking of fibrillar collagens and/or elastin have been associated with arterial stiffening with age and cardiovascular disease [10, 12– 16]. Carotid arteries express three main fibrillar collagens, with collagen-I>collagen-III>collagen-V in abundance [17, 18].

Cells sense changes in the stiffness of their microenvironment though a process called mechanotransduction, which typically involves the binding of ECM components to cell surface receptors and initiation of receptor-mediated signaling. Fibrillar collagens can bind to the discoidin domain receptor tyrosine kinases [19], but most studies have focused on the binding of fibrillar collagens to their receptors within the integrin family [20]. Integrin engagement leads to the recruitment of numerous signaling proteins and results in organization of the actin cytoskeleton, cellular stiffening and stiffness-mediated gene transcription [21–24]. The Rho and Rac nonreceptor GTPases are among the major intracellular signaling components regulated by changes in ECM stiffness. In turn, the stiffness-dependent activation of these GTPases has profound effects on the actin cytoskeleton, migration, proliferation and differentiation [25–34]. The third nonreceptor GTPase, Cdc42, also affects actin organization and motility [25, 35], but its role in SMC proliferation and differentiation is not well understood.

We previously reported that a stiffness-dependent activation of Rac, but not Rho, promotes SMC proliferation by stimulating G1 phase cyclin D1 induction and S phase entry [9, 36]. Consistent with these results, genetic depletion of SMC Rac1 strongly reduces cyclin D1 expression, proliferation, and neointimal formation after *in vivo* femoral artery injury [36]. However, the stiffness-dependent activation of Rac can also regulate SMC differentiation [31], which could then lead to changes in ECM production and arterial stiffness as described above. This scenario raised the possibility that the reduced SMC proliferation observed in injured Rac1-deficient arteries [36] might be secondary to potential Rac1 effects on SMC differentiation. To address this issue, we have now examined the levels of SMC differentiation markers and the major arterial collagens as well as elastin integrity in carotid artery sections of mice deficient for SMC Rac1. We also used pressure myography to compare carotid artery mechanics and contraction in the presence and absence of SMC Rac1.

## Results

### Smooth muscle-specific deletion of Rac1

We generated floxed Rac1 mice harboring a SMC-specific, tamoxifen-inducible Cre transgene (hereafter called Rac1^ff^;iCre mice) and treated the mice with vehicle (ethanol) or tamoxifen. Consistent with our previous work [36], Rac1 was strongly expressed in the carotid artery medial layer of Rac1^ff^;iCre mice treated with vehicle and much reduced in the carotid medial layers of mice that had received tamoxifen (Fig. 1A-B). We note an unexpected tamoxifen effect on adventitial Rac1, but the levels of Rac1 in the adventitia were low as compared to those in the media (Fig. 1A-B). Carotid artery morphology, as determined by H&E staining, appeared normal in both the vehicle and tamoxifen-treated mice (Fig. S1 top panels).

**Figure 1.**
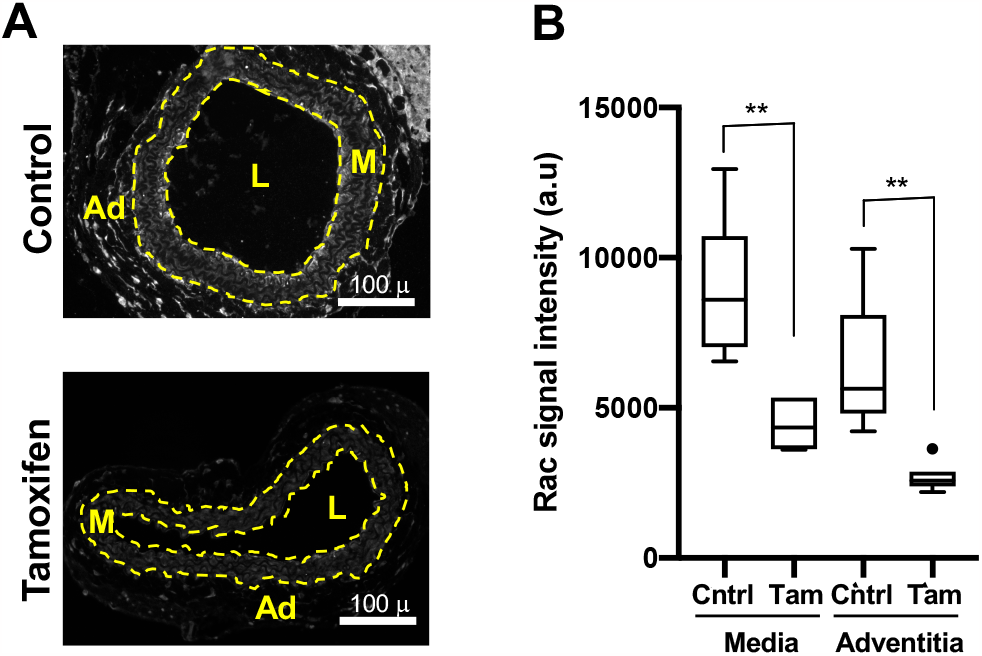
Deletion of smooth muscle Rac1. **(A)** Images of carotid artery cross-sections from Rac1^f/f^;iCre mice that had been treated with ethanol (vehicle; control) or tamoxifen. The sections were immunostained with an antibody to Rac1. Scale bars = 100 μm. The dashed lines in panel A approximate the positions of the external and internal elastic laminae, which separate the medial (M) and adventitial (Ad) layers. The lumen is indicated as “L”. **(B)** Immunostaining results, as shown in A, were quantified and graphed as box plots with Tukey whiskers; n = 6 per condition. Statistical significance was determined by Mann–Whitney tests, comparing control versus tamoxifen treatment within the medial and adventitial layers.

### Arterial mechanics

We performed biaxial inflation-extension tests on a pressure myograph to assess the effects of Rac1 deletion on arterial responses to increasing pressure. Axial arterial stiffness, as determined by the “*in vivo* stretch (IVS; see Methods and [37, 38]) and axial stretch-strain relationship, were not affected by the deletion of SMC Rac1 (Figs. 2A-B and S2). Similarly, changes in outer diameter (Fig. 2C), inner radius (Fig. 2D), wall thickness (Fig. 2E), or arterial stiffness (as determined by absence of right or left shifts in the stretch-strain curves; Fig. 2F) were not distinguishable in the carotid arteries containing or deficient for Rac1. Thus, deletion of SMC Rac1 did not affect these major parameters of arterial mechanics.

**Figure 2.**
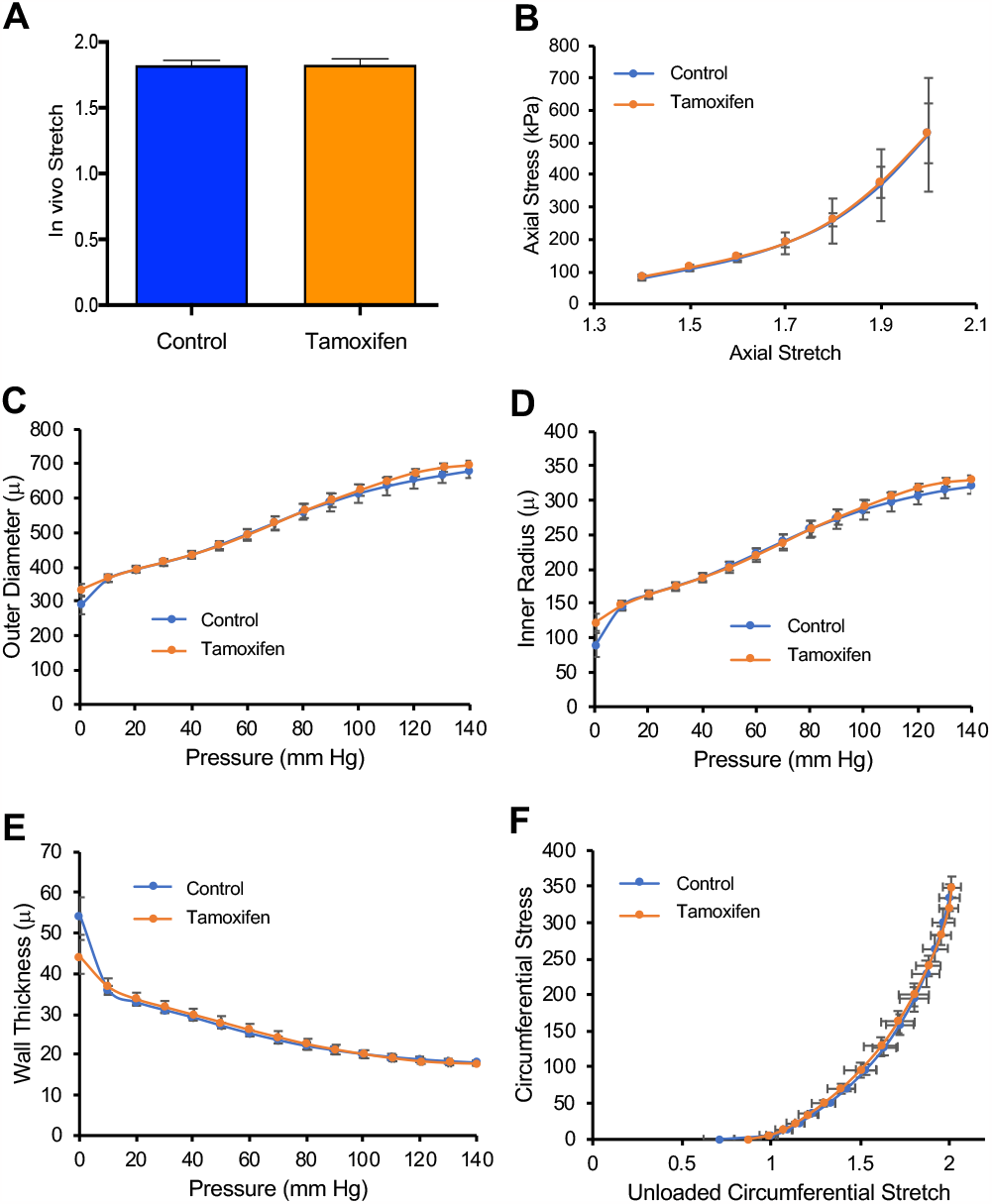
Deletion of SMC Rac1 does not affect passive arterial mechanics. Freshly isolated carotid arteries from control mice (n=6) or Rac1^f/f^;iCre mice treated with tamoxifen (n=7) were analyzed by pressure myography. To exclude potential effects of tamoxifen beyond activation of the Cre transgene, graphs for the control mice show combined results from Rac1^f/f^;iCre mice that had been treated with ethanol (vehicle; n=2) and Rac1^f/f^ mice (lacking the iCre transgene) that had been treated with tamoxifen (n=4). **(A)** *In vitro* stretch as determined from axial force-length tests. **(B)** Axial stress– stretch curves obtained at 80 mm Hg. **(C)** Changes in outer diameter with increasing pressure. **(D and E)** Changes in vessel inner radius and wall thickness, respectively, with pressure. **(F)** Circumferential stress–stretch curves with data points showing results at 0–140 mm Hg in increments of 10 mm. Results in panels A-B show means + SD, and results in panels C-F show means ± SE (see Methods). Statistical significance was determined by a Mann-Whitney test (panel A) or two-way ANOVAs (panels B-F) comparing control to tamoxifen. In F, statistical significance was tested for both stretch and stress.

### Fibrillar collagens, SMC differentiation and arterial contraction

The abundance of the arterial fibrillar collagens (I, III and V) as well as their expected preferential localization to the adventitial (collagen-I) or medial (collagen-III and -V) layers were similar in the vehicle- and tamoxifen-treated carotid arteries (Fig. 3A-B). Elastin integrity was also unaffected by tamoxifen treatment (Fig. S1 bottom panels).

**Figure 3.**
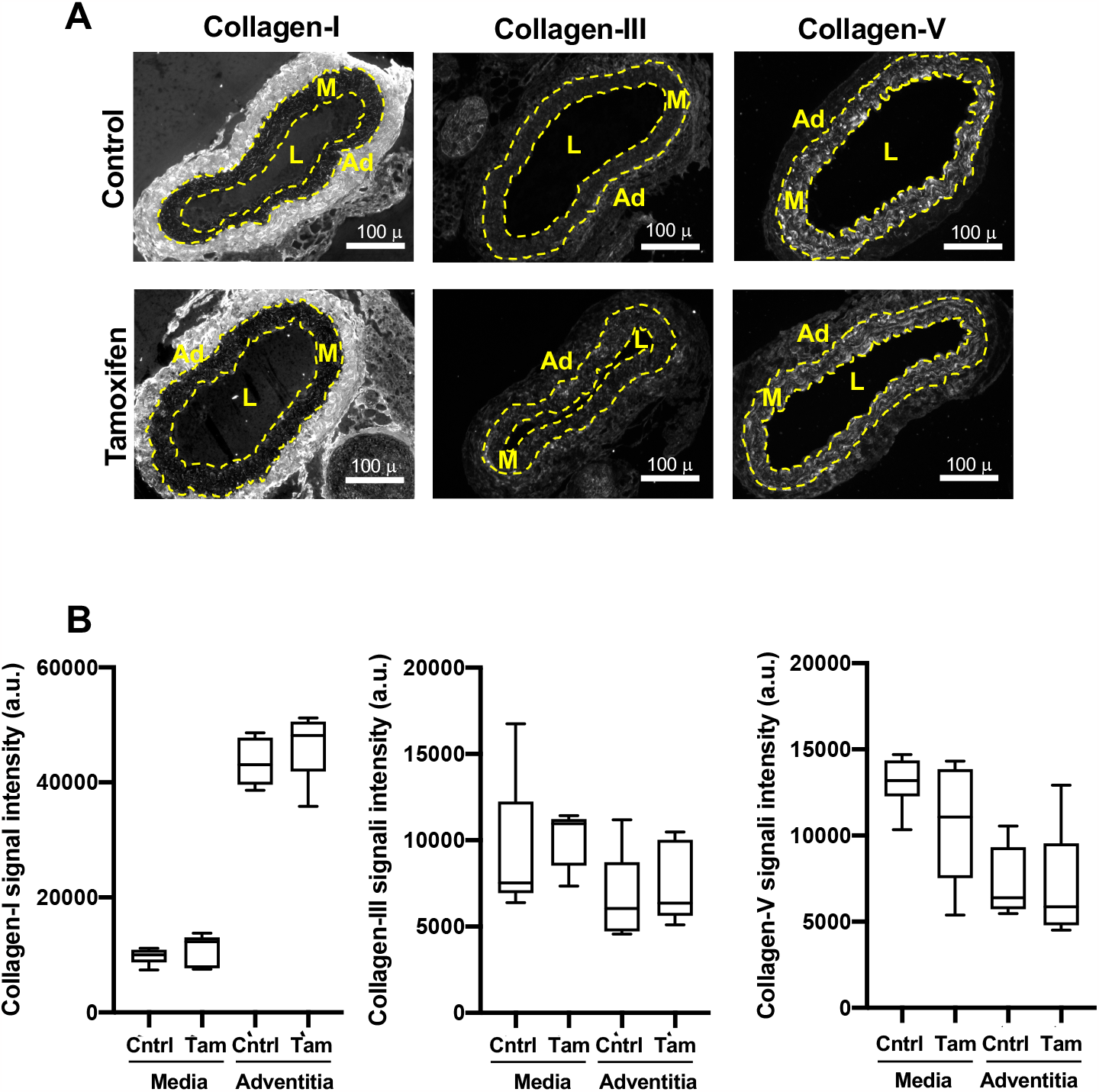
Arterial fibrillar collagen abundance is not affected by deletion of smooth muscle Rac1. **(A)** Images of carotid artery cross-sections from Rac1^f/f^;iCre mice that had been treated with ethanol (vehicle; control) or tamoxifen. The sections were stained with antibodies to collagen-I, collagen-III, or collagen-V. Scale bars = 100 μm. Dashed lines approximate the positions of the external and internal elastic laminae, which separate the medial (M) and adventitial (Ad) layers. The lumen is indicated as “L”. **(B)**Immunostaining results, as shown in A, were quantified and graphed as box plots with Tukey whiskers; n=6 per condition. Statistically significant differences were not detected when comparing (by Mann–Whitney tests) control versus tamoxifen treatment within the medial and adventitial layers.

The expression of medial SM-MHC trended lower in the carotid arteries of the Rac1-deficient mice, but the change was not statistically significant. The levels of medial α-SMA were not affected by Rac1 deficiency (Fig. 4A-B). Consistent with these results, contraction to KCl was not statistically different in the control and Rac1-deficient carotids (Fig. 4C).

**Figure 4.**
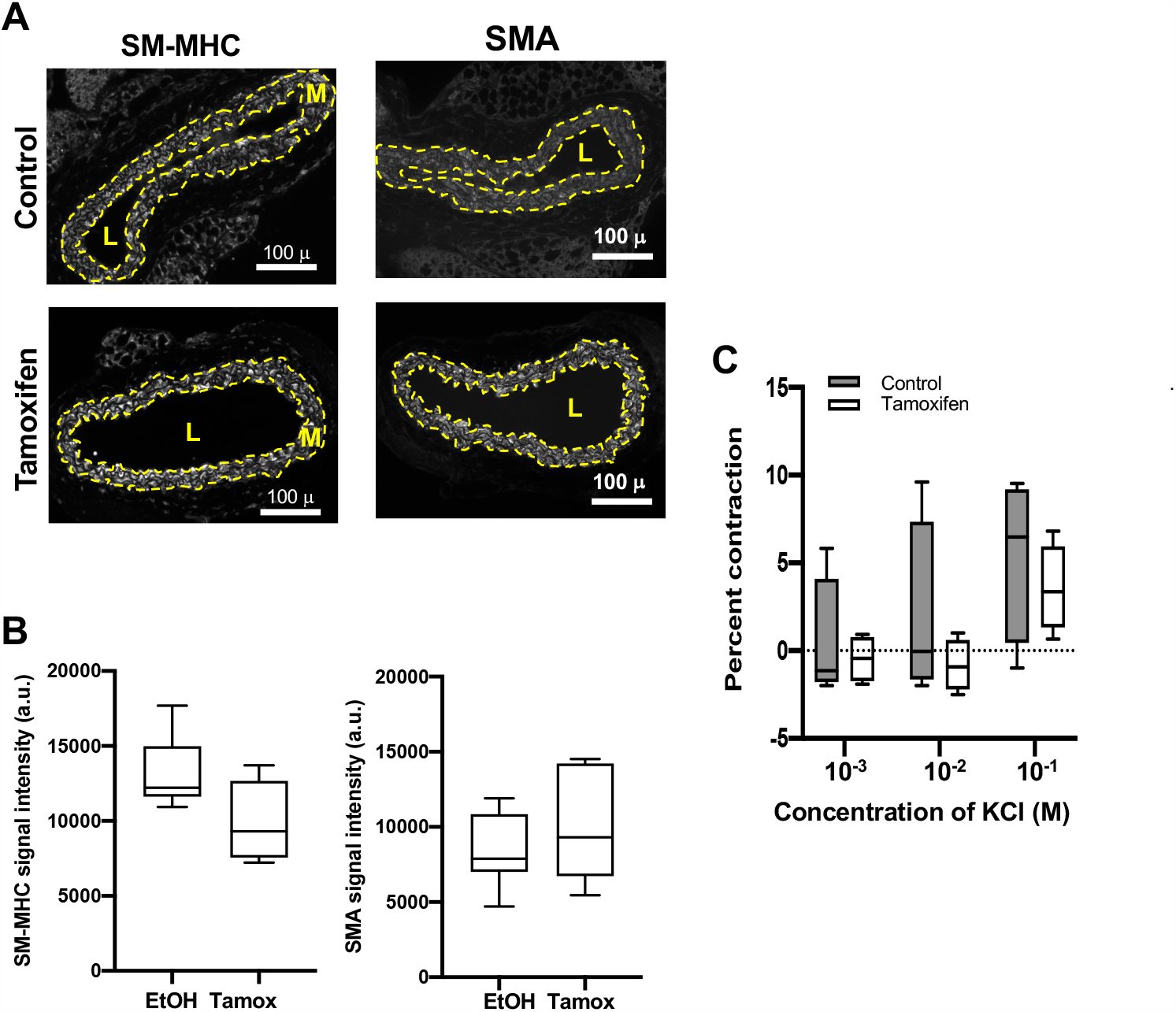
Minimal effect of *in vivo* Rac1 deletion on smooth muscle differentiation. **(A)** Representative images of carotid artery cross-sections from Rac1^f/f^;iCre mice treated with ethanol (vehicle; control) or tamoxifen. The sections were immunostained with an antibody to SM-MHC or α-SMA. Dashed lines approximate the positions of the external and internal elastic laminae, which encompass the medial layer (M). The lumen is indicated as “L”. Scale bars = 100 μm. **(B)** Immunostaining results for medial SM-MHC (n=6) and medial α-SMA (n=7) were quantified and graphed as box plots with Tukey whiskers. Statistical significance was tested using Mann–Whitney tests. **(C)** Contraction to KCl in isolated carotid arteries from mice containing Rac1 (control; n=4) or deficient in Rac1 (tamoxifen; n=4). To exclude potential effects of tamoxifen beyond activation of the Cre transgene, the contraction graph for the control mice show combined results from Rac1^f/f^;iCre mice that had been treated with ethanol (vehicle; n=2) and Rac1^f/f^ mice (lacking the iCre transgene) that had been treated with tamoxifen (n=2). The contraction seen in response to 10^-4^ M KCl was set to zero (dashed line in panel C) as this KCl concentration approximated that of the HBSS buffer. Statistical significance in C was tested using ANOVA, comparing contraction in the control vs. tamoxifen treated samples for each KCl concentration.

## Discussion

Several studies have shown that Rho activity controls the differentiation status of vascular SMCs by promoting the differentiated (contractile) phenotype. However, Talwar et al. [31] recently reported that Rac acts together with Rho to maintain SMCs in the contractile state. We therefore wondered if arterial depletion of Rac1 (but not Rho) might be sufficient to affect SMC differentiation and its consequent effects on ECM remodeling and arterial stiffness.

The results here show that Rac1-containing and -deficient carotid arteries similarly express SMC differentiation markers and the major arterial fibrillar collagens. Moreover, Rac1 deficiency affects neither the integrity of medial elastin nor normal arterial mechanics as judged by pressure myography. Similarly, the control and Rac1-deficient carotid arteries did not show a statistically significant difference in their contractile response to KCl. Thus, *in vivo* Rac1 deficiency in itself does not appear to alter the basal, contractile state of arterial SMCs. These findings are notably different from those we recently described in carotid arteries deficient for focal adhesion kinase (FAK), as medial FAK deletion was sufficient to reduce arterial contraction to KCl [37]. Since FAK activity is upstream of Rac-GTP activation [9, 36], FAK effectors beyond Rac must be required to regulate arterial contraction.

These results also inform our understanding the effect of Rac1 on SMC proliferation. If Rac1 deficiency had altered SMC differentiation, then the altered basal differentiation state could have contributed to the reduced SMC proliferation observed in Rac1-deficient femoral arteries after vascular injury [36]. However, this was not the case. While we cannot formally exclude potentially distinct effects of Rac1 in femoral [9] versus carotid (this report) SMCs, our combineπd results generally support the idea that the reduced proliferation of Rac1 deficient arteries in response to vascular injury reflects the stimulatory effect of Rac1 signaling to the cell cycle and is not a secondary consequence of Rac1 effects on SMC differentiation or arterial stiffness.

## Methods

### Mice and artery isolation

Mice on the C57BL/6 background harboring floxed Rac1 genes and a tamoxifen-inducible, SMC-specific Cre transgene were generated as described [36]. Since the SMC-specific Cre transgene is on the Y-chromosome [39], all experiments were performed on male mice. The mice were fed a chow diet *ad libitum*, aged to 3-4 months, treated with ethanol (vehicle) or tamoxifen (Sigma Aldrich) for 5 days as described [36], allowed to recover for 1–2 weeks, and sacrificed by CO_2_ asphyxiation at 4-5 months. The right carotid artery was immediately removed, stripped of most fat, and used for pressure myography. The left carotid artery was perfused *in situ* with PBS, removed, cleaned in PBS and fixed in Prefer (Anatech Ltd.) for paraffin-embedding, sectioning and immunostaining. Animal protocols were approved by the University of Pennsylvania Institutional Animal Care and Use Committee.

### Pressure myography

Arterial mechanics and the contractile response to KCl were determined *ex vivo* on a DMT 114P pressure myography with force transducer largely as described [37, 38, 40]. Isolated right carotid arteries (see above) were secured to 380 µm (outer diameter) cannulas using silk sutures, and submerged in 5-ml of calcium-containing Hanks Balanced Salt Solution (HBSS; Corning Life Sciences). This closed system was checked for leaks by pressurizing the vessel to 30-mm Hg using HBSS and ensuring that there was no pressure loss through the system. The arteries were also visualized by light microscopy, and the axial length (where the artery transitioned from being bent to straight) was recorded as the unstretched vessel length (UVL). Arteries were then brought to a stretch of 1.8 and pressurized to 100 mm Hg for 15 min with calcium-containing HBSS. The arteries were preconditioned by pressurization three times from 0–140 mm Hg. Unloaded (unstretched and unpressurized) vessel wall thickness and outer diameter were measured in multiple sections after preconditioning and averaged with measurements after axial testing for post-test data analysis. *In vivo* stretch (IVS) was determined using force-length tests as described [37, 38], and the intersection of the force-stretch curves was defined as the IVS. Loaded inner radius and wall thickness were determined from pressure-outer diameter tests with samples at their IVS and pressurized in 10-mm Hg steps from 0-140 mm Hg (30-sec per step) before returning the artery to 0 mm Hg. The validity of IVS determinations was assessed by measuring axial force through the circumferential tests; we excluded samples where axial force varied from the mean by >25% with pressure. Raw measurements of the intraluminal pressure, force transducer readings, and video-tracked outer diameters were converted into stress-stretch curves as described [37, 38]. Vessel wall thickness was calculated in the post-test analysis based on the sample being incompressible [38].

To measure the contractile response to KCl, freshly isolated left carotid arteries were cleaned of excess fat, mounted on the myograph, and incubated in 5 ml of HBSS as described above. The closed system was checked for leaks, returned to 0 mm Hg, and the UVL was determined, also as described above. Mounted vessels were preconditioned by stretching them axially to 1.15 times their UVL at 40 mm Hg for 15 min and then 1.3 times their UVL at 60 mm Hg for 15 min. Vessels were then brought to their IVS (approximately 1.8) and pressurized to 100 mm Hg. The vessel was allowed to equilibrate for 5 min. After baseline diameter measurements were taken, increasing concentrations of KCl (10^-4^ M to 10^-1^ M in 5 ml HBSS) were added to the bath sequentially by first removing the preceding bath and adding the new dilution. Each dilution was allowed to reach a constriction plateau (∼5-10 min) before outer diameter measurements were recorded and the next dilution added. KCl was rinsed out with HBSS until >95% of the original outer diameter was attained. For all treatments, outer diameters were recorded in real time at each KCl concentration using MyoVIEW software (DMT) and an inline tracking camera (Imaging Source).

### Immunofluorescence staining

Paraffin sections of isolated carotid arteries were prepared and immunostained for collagen-I, collagen-III, collagen-V, α-SMA, and SM-MHC as described [37, 40, 41]. Rac immunostaining used anti-Rac1 (catalog # 610650; BD Transduction Laboratories), typically with a ∼100-fold primary antibody dilution and an Alexa Fluor secondary antibody (ThermoFisher). Results were visualized with a Nikon Eclipse 80i microscope with a QI-Click Qimaging camera. Carotid arteries were imaged at 20 x magnification. Images were quantified using Fiji. The media of each section was traced using the polygon drawing tool. Raw integrated intensity was divided by the area of the outlined media to yield values for relative fluorescence intensity. This procedure was repeated for the adventitial layer.

### Statistical Analysis

The statistical tests were performed using Prism software (GraphPad). For the pressure myography experiments, *in vivo* axial stretch and the axial stress-stretch curves were determined from single measurements; graphs show mean ± SD. Pressure-mediated changes in outer diameter, inner radius, wall thickness, and circumferential stretch-stress results were determined from multiple measurements (see [38]); graphs show mean ± SE. Two-tailed Mann-Whitney tests were used to compare two datasets, and 2-way ANOVAs were used for multiple comparisons. Statistical significance for all graphs is demarcated by *(p<0.05), **(p<0.01), ***(p<0.001). Box plots show Tukey whiskers.

## Acknowledgements

We thank V. Tybulewicz (National Institute of Medical Research, London) for the floxed Rac1 mice, and S. Offermanns (Max Planck Institute for Heart and Lung Research) for the smooth muscle–specific, inducible Cre mouse. This work was supported by NIH grant HL137232 to RKA.

---

**Figure S1.**
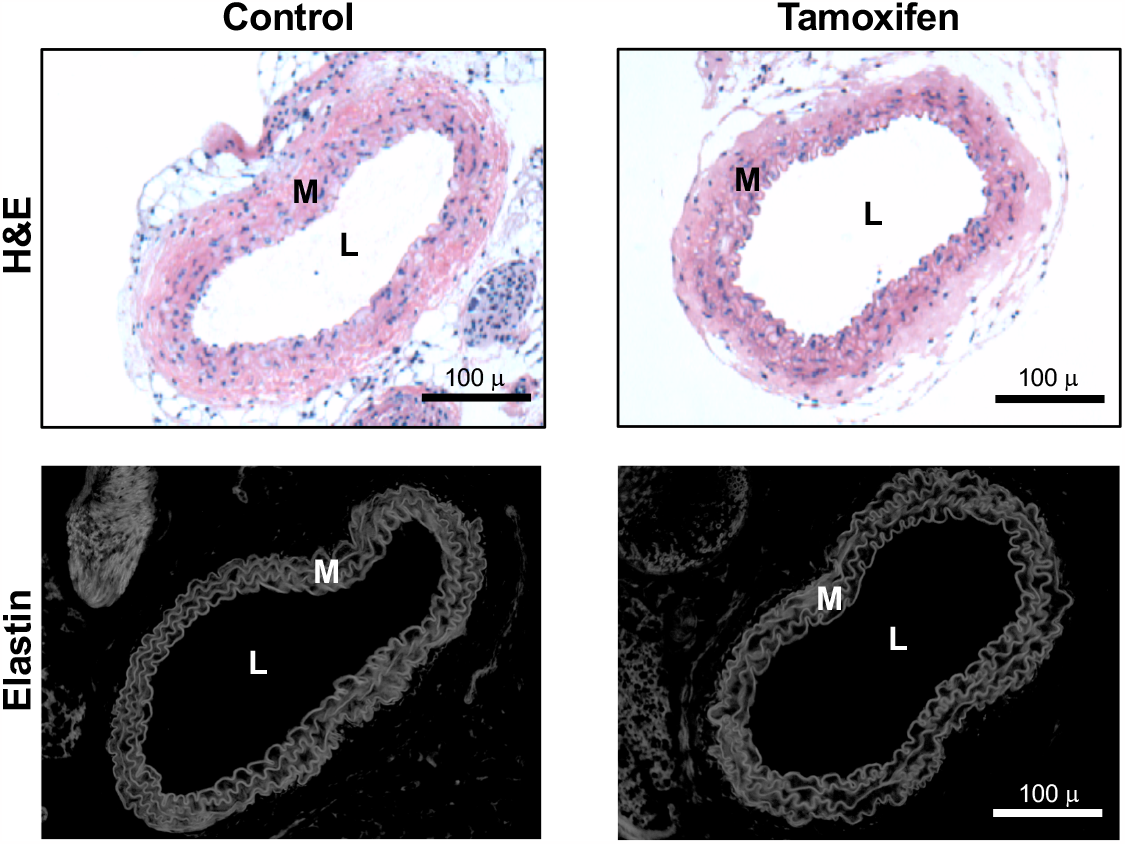
Histological characterization of Rac1^f/f^;iCre mice. H&E staining (top panels) and autofluorescence images of medial elastin (bottom panels) in isolated carotid arteries of Rac1^f/f^;iCre mice treated with ethanol (vehicle; control) or tamoxifen. Images are representative of 7 control and 7 tamoxifen-treated mice. The media and lumen are indicated as “M” and “L”, respectively.

**Figure S2.**
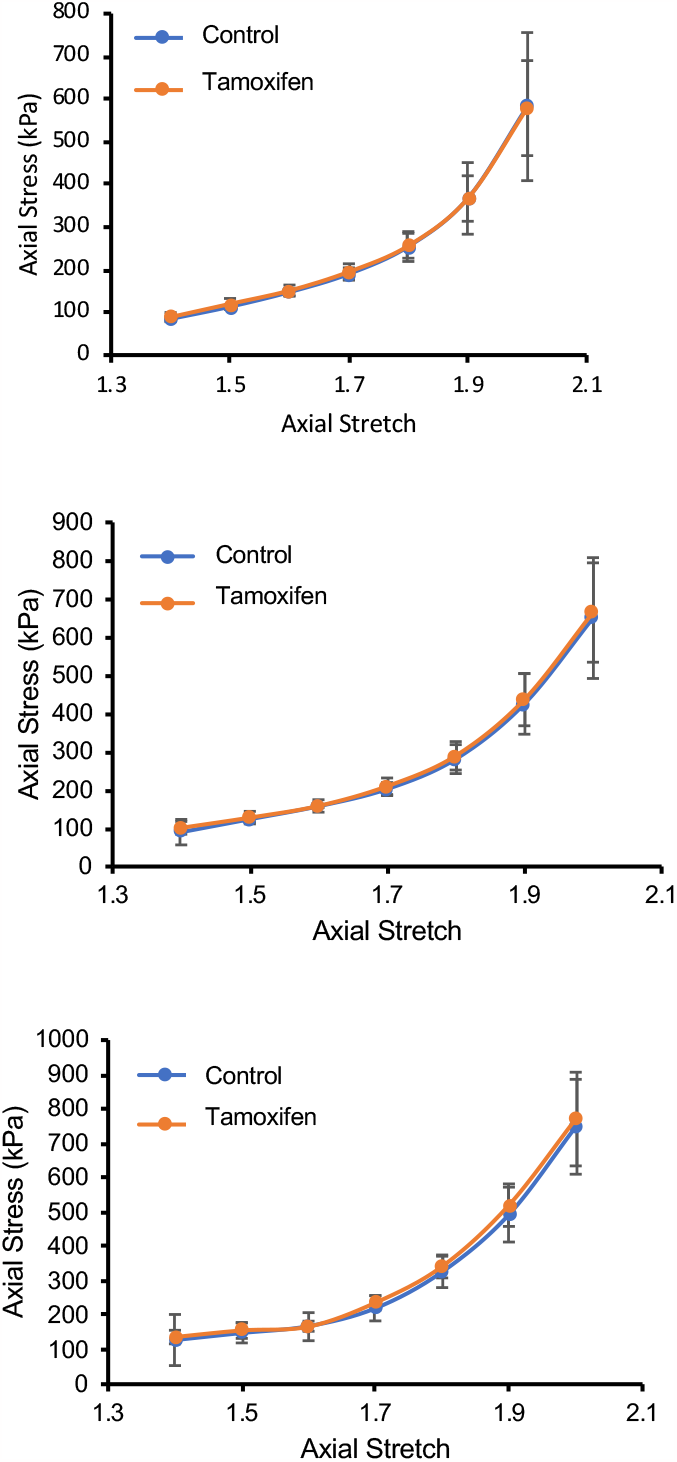
Additional axial stress-stretch curves. The figure compiles axial stretch-stress curves generated at 100 (top), 120 (middle) and 140 (bottom) mm Hg for the control and Rac-deficient carotid arteries shown in Figure 2.

